# Degradation of ribosomal RNA during *Plasmodium falciparum* gametocytogenesis

**DOI:** 10.1101/2025.02.12.637867

**Authors:** Janne Grünebast, Ritwik Singhal, Robin Bromley, Sachie Kanatani, Kaylee Watson, Franck Dumetz, Tales Vicari Pascini, Abhai Tripathi, Julie C Dunning Hotopp, Photini Sinnis, Manuel Llinás, David Serre

## Abstract

The life cycle of *Plasmodium falciparum* is characterized by complex regulatory changes that allow adaptation of the parasites to different environmental conditions, which are especially pronounced during transmission between the mammalian host and the insect vector. Previous studies have shown that *P. falciparum* uses three types of ribosomal RNAs (rRNA A-, S1- and S2-types) at different stages of its life cycle. We used Oxford Nanopore Technologies (ONT) direct RNA sequencing to investigate the dynamics of rRNA usage throughout the parasite’s intraerythrocytic development, as well as in salivary gland sporozoites. Our study revealed a preponderance of A-type rRNAs during the intraerythrocytic cycle and gametocytogenesis, while S-type rRNAs slowly increase in abundance in mosquito stages starting three days post infection. Salivary gland sporozoites showed an even proportion of all rRNA types. By examining the length distributions of rRNA molecules, we detected an extensive and specific degradation of rRNAs during gametocytogenesis, starting in stage II gametocytes and continuing until the final stages of gametocyte development. We hypothesize that rRNA degradation may be linked to the global translational repression and metabolic quiescence described in stage V gametocytes, similar to mechanisms observed in bacterial and eukaryotic stress responses.

## Introduction

Malaria remains one of the world’s major global health threats, with 263 million new cases and over half a million deaths reported in 2023 alone [1]. The disease is endemic to 83 countries, with 94% of cases occurring in Africa, where it mainly affects sub-Saharan African children under the age of five and remains one of the leading causes of death [2]. The primary causative agent of malaria in Africa is *Plasmodium falciparum*, a unicellular protozoan parasite from the *Plasmodium* genus. *Plasmodium* parasites have a complex life cycle and go through several life cycle stages in very different environmental conditions in both the mosquito and the mammalian hosts [3]. Transmission occurs when a female *Anopheles* mosquito injects sporozoites into a human host. These sporozoites travel in the bloodstream to the liver, where they invade hepatocytes and undergo extensive asexual replication. Upon rupture, hepatic schizonts release tens of thousands of merozoites into the bloodstream, initiating the intraerythrocytic stage of the life cycle. In red blood cells, the parasites undergo cycles of replication, transitioning through ring, trophozoite, and schizont stages, before rupturing to release more merozoites. This cycle of replication and release causes all of the clinical symptoms associated with malaria. A subset of blood-stage parasites differentiates into sexual forms, the gametocytes, in a 12-16-day process in *P. falciparum* that involves five distinct morphological stages (I-V). Stage V gametocytes are taken up during a blood meal and establish infection in the mosquito host [4]. Within the mosquito, gametocytes rapidly differentiate into microgametes and macrogametes, which fuse to form a zygote. The zygote then develops into an ookinete, which crosses the mosquito midgut wall to form an oocyst, where sporozoites develop and eventually migrate to the mosquito salivary glands, ready for transmission to a new host [5].

This complex life cycle is orchestrated by gene expression changes that are tightly regulated throughout malaria parasite development. While the speed of development varies among *Plasmodium* species and may be influenced by environmental factors, *Plasmodium* parasites typically progress continuously throughout their life cycle, with two critical exceptions when the parasites pause their development [6, 7]. Mature stage V gametocytes in the bloodstream can remain infectious for weeks until they are picked up by a mosquito, while sporozoites reside in the mosquito salivary glands waiting to be injected into a human host [5]. During these two stages, the parasites reduce their metabolic activity but need to be poised to rapidly re-initiate development upon transmission. For example, gametogenesis in the mosquito midgut is completed within minutes [8–10] while sporozoites become motile upon injection in the dermis [11–13]. One of the mechanisms underlying these developmental arrests and rapid reactivation upon transmission is the translational repression of mRNAs that allows the parasite to rapidly synthesize proteins from stored mRNAs after transmission [14–20].

Ribosomal RNAs (rRNAs) are essential for translating mRNAs into proteins, and many eukaryotes have hundreds of copies of these genes. By contrast, *Plasmodium* parasites only have four to five copies, with slightly different rRNA sequences expressed at different times of the life cycle [21–23]. *P. falciparum* has three types of rRNAs (A-type, S1-type, and S2-type) encoded by five gene clusters, each containing a copy of the 18S, 5.5S and 28S rRNA genes arranged in tandem and transcribed polycistronically by RNA polymerase I [24, 25]. The nucleotide sequences of the two A-type rRNAs, found on chromosomes 5 and 7, are nearly identical but differ from the S1-type rRNA sequence on chromosome 1 and from the S2-type rRNA sequences on chromosomes 11 and 13. Although the nucleotide sequences for the two S2-type rRNA are almost identical, they are more variable, particularly in the 28S rRNA gene, compared to the S1-type rRNA sequence [25]. The main structural differences between the A- and S-type 28S rRNAs were identified at the GTPase site and are conserved across *Plasmodium* species [26, 27]. These different rRNA types may allow *Plasmodium* parasites to assemble ribosomes with distinct compositions, facilitating selective regulation of specific mRNAs in response to the varying environments of its human and mosquito hosts, similar to what has been described in bacteria and other eukaryotic organisms [28].

Studies of rRNA abundance are challenging in *Plasmodium* due to the high similarity among the different rRNA genes and the varying amount of RNA and rRNA present at different developmental stages. A-type rRNAs are the most abundant rRNAs during the blood (and liver) stages, where they support the rapid protein synthesis necessary for survival in the nutrient-rich human host [22, 29, 30]. These A-type rRNAs appear to be primarily transcribed during the ring stage and the nucleolus, the ribosome factory, disappears during replication of blood-stage parasites [31]. In the mosquito, the abundance of A-type rRNAs seem to gradually decrease, possibly due to cleavage of existing molecules, while S2-type rRNAs increase throughout sporozoite development [22]. The S1-type rRNAs appear to be transcribed at low levels throughout the entire life cycle [30] but do not seem to predominate in abundance at any point.

Here, we used highly specific qRT-PCR and direct RNA sequencing using Oxford Nanopore Technology (ONT) to analyze RNAs collected throughout i) the asexual life cycle and stage V gametocytes from *P. falciparum* NF54 parasites and from ii) a tightly synchronized gametocytes time course of the Southeast Asian Dd2 strain, and rigorously characterized the expression levels of the different *P. falciparum* rRNAs and their variations.

## Methods

### *P. falciparum* cultures and RNA isolation

To evaluate the expression of rRNAs throughout development, we analyzed RNA collected from two strains of *P. falciparum* (NF54 and Dd2) using different culture protocols. Asexual parasites and stage V gametocytes were collected from the NF54 strain at the University of Maryland School of Medicine (UMSOM) while asexual parasites and gametocytes through their development were collected from the Dd2 strain at The Pennsylvania State University (PSU). In addition, stage V gametocytes and sporozoites were collected from the NF54 strain at Johns Hopkins School of Public Health (JHSPH) and ookinetes and oocysts from the NF54 strain collected at Sanaria after infection of *Anopheles stephensi*. We describe below the detailed culture and isolation protocols utilized for each sample.

### NF54 culture and isolation protocols (UMSOM)

*P. falciparum* NF54 parasites were cultured at the University of Maryland School of Medicine at 4% hematocrit with parasitemia below 6%, in incomplete RPMI media containing 25 mM HEPES and L-glutamine (Corning Inc. Life Sciences) (supplemented with 0.24 % (w/v) sodium bicarbonate, 0.367 mM hypoxanthine and 25 µg/ml gentamycin) plus 0.25% (w/v) AlbuMAX® and 5% heat-inactivated human serum (Grifols Bio Supplies).

For asexual stages (rings, trophozoites, and schizonts), the culture was synchronized with sorbitol and the different stages collected at different time points, after confirmation by microscopy. Asexual parasites were lyzed with saponin, resuspended in 500 µl of TRIzol^®^ reagent (Thermo Fisher Scientific) and stored at −80°C until RNA isolation.

For collecting gametocytes, we used the “crash” method [32]. Briefly, when the culture reached 3–5% ring-stage parasitemia (day 0), we synchronized parasites using 5% sorbitol. The following day (day 1), the spent media was carefully removed, and fresh, pre-warmed incomplete media with 10% human serum was added. A Giemsa-stained smear was prepared to confirm the presence of ring-stage parasites. On day 2, when rings were observed, the culture was diluted to 0.1% rings. The media was changed daily without adding additional red blood cells until day 16 when stage V gametocytes were harvested. Smears were continued daily for gametocytogenesis. A multilayer Percoll gradient consisting of 80%, 65%, 50%, and 35% Percoll layers was prepared, with the parasites resuspended in incomplete media layered on top. Gametocytes were then collected from the 35/50% interface, lyzed with saponin, and resuspended in 500 µl of TRIzol^®^ reagent (Thermo Fisher Scientific).

Trizol samples were thawed on ice and, after incubating at room temperature for 5 minutes, 100 µl of chloroform was added for 500 µl TRIzol^®^ reagent (Thermo Fisher Scientific). The samples were then vortexed and centrifuged at 12,100 rpm for 5 minutes at 4°C to separate the phases. The aqueous phase was carefully transferred to a new tube, mixed with an equal volume of chloroform, and centrifuged again. The RNA from the aqueous phase was precipitated by adding isopropanol and GlycoBlue, incubating at room temperature for 10 minutes, and centrifugation at 12,100 rpm for 30 minutes at 4°C. The resulting RNA pellet was washed with fresh 75% ethanol, air-dried briefly, and dissolved in 20 µl of DEPC-treated water. RNA samples were immediately frozen and stored at −80°C to maintain integrity.

### Dd2 culture and RNA isolation protocols (PSU)

*P. falciparum* Dd2 parasites cultured at the Pennsylvania State University at 3% hematocrit were synchronized twice using a 5% sorbitol solution. At 20-24 hours post-invasion (hpi), parasites previously cultured in incomplete RPMI media with L-glutamine (Corning Inc. Life Sciences) (supplemented with 25 mM HEPES (pH 7.4), 0.2 % (w/v) sodium bicarbonate, 0.1 mM hypoxanthine and 50 μg/ml gentamycin) plus 0.25% AlbuMAX ® (complete media), were diluted to 1.5-2% parasitemia in 25-50 mL of mFA (minimal fatty acid) media: Incomplete RPMI media, 0.39% fatty-acid free BSA (Sigma-Aldrich), 30 uM Oleic Acid (Sigma-Aldrich), 30 uM Palmitic Acid (Sigma-Aldrich). On day 1, 20-22 hours post-induction, the media was replaced with complete RPMI media. From day 2 onwards, 20 U/mL of Heparin was added to the complete media. Asexual stage parasites were harvested at 10, 20, 30, and 40 hpi, centrifuged and stored in TRIzol^®^ reagent with Trizol added at three times the red blood cell (RBC) pellet volume (∼10 ml) (Thermo Fisher Scientific) for RNA isolation. Similarly, sexually committed rings were collected at 30 hours post mFA induction followed by gametocytes at days 2, 4, 6, 8, 12, and 16 post-induction [33].

After storage at −80°C, samples were thawed, and 2 ml of chloroform was added per 10 ml of Trizol. Following centrifugation at 3,000 rpm for 10 minutes at 4°C, the clear supernatant (upper phase) was transferred to a new tube, avoiding contamination. Sodium acetate (3 M, pH 5.5) and an equal volume of cold isopropanol were added, and RNA was precipitated overnight at −20°C. On the following day, samples were centrifuged at 9,000 rpm for 1 hour at 4°C to form RNA pellets, which were washed with cold 70% ethanol and centrifuged again. The RNA pellets were dried and resuspended in nuclease-free water. Equal volume of 7.5M Lithium Chloride solution was added to the samples which were then incubated overnight at 4°C. On the following day, samples were centrifuged at maximum speed for 35 minutes at 4°C using a microcentrifuge. The RNA pellet was washed twice with 1 ml of chilled 70% ethanol, centrifuging for 10 minutes between each wash. After the final ethanol wash, the pellets were dried and resuspended in 30-100 µl of nuclease-free water for downstream analysis. RNA was stored at −80°C.

### Generation of gametocytes and sporozoites (JHSPH)

Low passage *P. falciparum* NF54 blood stage cultures at Johns Hopkins School of Public Health were maintained *in vitro* in O^+^ erythrocytes at a 4% hematocrit in RPMI 1640 (Corning Inc. Life Sciences) supplemented with 2.1 mM L-glutamine, 25 mM HEPES, 0.72 mM hypoxanthine (Sigma-Aldrich), 0.21% w/v sodium bicarbonate (Sigma-Aldrich), and 10% v/v heat-inactivated human serum (Interstate blood bank). Cultures were maintained at 37°C in a glass candle jar. Gametocyte cultures were initiated at 0.5% asexual blood stage parasitemia at 4% hematocrit and maintained by changing the media daily for 17 days without the addition of fresh blood to promote gametocytogenesis as described earlier [34]. Erythrocytes used for the study were obtained from healthy donors under a Johns Hopkins Institutional Review Board-approved protocol and were provided without identifiers to the laboratories. Gametocytes were obtained as described above for *P. falciparum* NF54 parasites (“crash” culture) and purified with a Percoll gradient, saponin lyzed, and resuspended in 500 µl Trizol reagent. RNA isolation followed the protocol from the NF54 samples in the section above (replicate #2).

Adult *Anopheles stephensi* mosquitoes (3 to 5 days post emergence) were allowed to feed through a glass membrane feeder for up to 30 minutes on *P. falciparum* NF54 cultures with gametocytemia of 0.3% in 40% hematocrit containing fresh O^+^ human serum and O^+^ erythrocytes. Infected mosquitoes were maintained at 25°C and 80% humidity and were provided with 10% w/v sucrose solution for 19 days. Salivary glands of 100 mosquitoes were dissected in 1X PBS, manually homogenized and sporozoites were spun down at 14,000 g for 5 minutes. The pellet was resuspended in 500 µl of Tri Reagent (Molecular research center), vortexed for 1 minute, and stored at −80°C until RNA isolation (following the NF54 RNA isolation protocol).

### RNA isolation from parasites after mosquito infection (Sanaria)

Human O^+^ erythrocytes were obtained weekly from Vitalant® (Memphis, TN). The erythrocytes were washed upon arrival with 0.2-µM filtered RPMI 1640 medium (containing 25 mM HEPES and 50 µg/ml hypoxanthine; KD Medical, Columbia, MD) and stored at 50% hematocrit in a 4°C refrigerator for use within a week from processing. Asynchronous cultures of the *P. falciparum* NF54 [35] were kept at 10 ml volumes in T25 vented flasks (Corning Inc. Life Sciences) at 5% hematocrit with complete RPMI medium (RPMI 1640 medium supplemented with 10 mg/l gentamicin, 0.23% sodium bicarbonate, and 10% O^+^ pooled human serum (Vitalant^®^)). Cultures were incubated at 37°C with a gas mixture (90% N_2_, 5% O_2_, and 5% CO_2_). Medium was changed daily. Parasitemia was monitored by methanol-fixed thin blood films stained for 15 minutes with 20% Giemsa solution (Sigma-Aldrich) and maintained between 0.5 and 9% parasitemia. *P. falciparum* gametocytes were induced *in vitro* [36]. Briefly, cultures of *P. falciparum* NF54 parasites were initiated as mixed stages in T75 flasks at 0.5% parasitemia and 5% hematocrit in the complete RPMI medium described above. Cultures were maintained with daily medium changes. Media changes were done on top of a slide warmer unit at 37°C. When stage V gametocytemia was prevalent at > 0.5% (days 14–16), the culture was collected for mosquito feeding. A stage V gametocyte sample (replicate #3) at the time point of infection was harvested by centrifugation for 5 minutes at 3000 rpm and the pellet was stored in Trizol for RNA isolation.

*Anopheles stephensi* SDA500 [37] were reared at Sanaria Inc. insectary at 27°C and 80% relative humidity, with a 12 h/12 h light/dark cycle. Larvae were reared in trays with dechlorinated water and fed brewer’s yeast and fish food (Aqueon^®)^, while adults were maintained with cotton pads soaked in water and sugar cubes (Domino^®^ Sugar, Baltimore). For colony maintenance and egg production, the females were fed with human whole blood (Vitalant^®^) using artificial membrane feeders. Three-to-four day-old *An. stephensi* females were heat-selected and transferred into a one-gallon paper container (Science Supplies WLE Corp.) and blood-fed with whole blood containing *P. falciparum* gametocyte culture. All operations with infected live mosquitoes were performed inside a secure, triple-screened insectary according to Sanaria’s SOPs. After feeding, the containers of mosquitoes were maintained inside the incubator (Sanyo^®^) at 26°C, 80% humidity, and 12 h/12 h light/dark cycle). The mosquitoes were provided a water-soaked cotton wool ball and sugar cubes daily. Mosquito samples were collected at 2 h, 24 h and 3 days post-infection (dpi). For each time point, females were aspirated from containers, knock-down on freezer for 12 minutes, washed on ethanol for 2 minutes, and then placed on PBS 1X for dissection. Midguts from ten females were dissected on PBS, pooled on an Eppendorf tube containing TRIzol^®^ reagent (Thermo Fisher Scientific) and homogenized using an automatic tissue pestle. Tubes were stored at −80°C until the moment of RNA extraction. RNA isolation followed the protocol for the NF54 culture.

### Oxford Nanopore direct RNA sequencing

500 ng of total RNA was used for preparing Direct RNA sequencing library for each sample (except after *in vitro* polyadenylation when 150 ng of RNA was used, see below) using the SQK-RNA002 kit (Oxford Nanopore Technologies) and following the manufacturer’s protocol. Briefly, RNA samples were ligated to an RT adapter using T4 DNA Ligase, followed by reverse transcription with Induro Reverse Transcriptase (New England Biolabs). The resulting cDNA-RNA hybrids were purified using Agencourt RNAClean XP beads (Beckman Coulter) and eluted in nuclease-free water. After a second ligation step and clean-up, the library was loaded onto a MinION flow cell (R9.4.1) for direct RNA sequencing using the MinKNOW software (Oxford Nanopore Technologies). Base-calling options were turned off and the sequencing run continued for up to 72 hours or until the flow cell was fully depleted.

### *In vitro* polyadenylation

*In vitro* polyadenylation of 150 ng RNA was performed for NF54 trophozoites and stage V gametocytes using *E. coli* Poly(A) Polymerase (New England Biolabs) according to the manufacturer’s protocol. The reaction was incubated at 37°C for 75 sec. To stop the reaction, EDTA was added to a final concentration of 10 mM, and the reaction was directly used for library preparation using the Direct RNA Sequencing Kit as described above.

### Analyses of rRNAs using qRT-PCR

For RT-qPCR experiments, we first ensure that contaminating genomic DNA (gDNA) was removed from the RNA extracts using DNA-*free*^TM^Kit (Thermo Fisher Scientific) following the manufacturer’s protocol. Briefly, each RNA sample was treated with DNase I by adding 0.1 volume of 10X DNase I buffer and 1 µl of rDNase I, followed by incubation at 37°C for 30 minutes. After incubation 2 µl (or 0.1 volume) resuspended DNase Inactivation Reagent was added to each sample and incubated at room temperature for 2 minutes with intermittent mixing. The samples were centrifuged at 10,200 rpm for 1.5 minutes, and the RNA-containing supernatant was transferred to a fresh tube, avoiding disturbance of the DNase Inactivation Reagent pellet. RNA was tested for gDNA contamination using primers spanning the intron-exon region of HSP90 (see Supplementary Table S1 for the primer sequences). A gDNA control extracted using the Dneasy® Blood & Tissue Kit (Qiagen) was used as a positive control and for all RT-qPCR experiments.

For cDNA synthesis, the SuperScript^TM^ II Reverse Transcriptase (Invitrogen) Kit was used. Briefly, 10 µl of RNA (1 ng/µl) was combined with 0.5 µl of random hexamers (Promega), 1 µl of 10 mM dNTP mix, and 0.5 µl of nuclease-free water. The mixture was heated to 65°C for 5 minutes, chilled on ice, and briefly centrifuged. 4 µl of First-Strand Buffer, 2 µl of 0.1 M DTT, and 1 µl of Superase Inhibitor were then added and gently mixed. The reaction was incubated at 25°C for 2 minutes, followed by the addition of 1 µl of SuperScript II Reverse Transcriptase. The mixture was incubated at 25°C for 10 minutes, then at 42°C for 50 minutes. The reaction was inactivated by heating to 70°C for 15 minutes. For the qPCR reaction, 5 µl of 2x SYBR Green mix was combined with 0.4 µl of forward primer (10 µM), 0.4 µl of reverse primer (10 µM), and 3.2 µl of nuclease-free water. A total of 9 µl of this qPCR master mix was added to 1 µl of the cDNA template. The qPCR was performed using a 3-step protocol: an initial denaturation at 95°C for 3 minutes, followed by 40 cycles of 95°C for 10 seconds, 55°C for 10 seconds, and 72°C for 10 seconds.

Primer specificity was tested using three different DNA constructs (IDT minigenes) made of the sequence of the first 700 bp from the 28S rRNA from A-type (PF3D7_0532000), S1-type (PF3D7_0112700), and S2-type (PF3D7_1148640), respectively. The three gDNA controls (10 ng, 1 ng, 0.1 ng) and the three different rRNA type constructs were included in all qPCR runs. qPCR results were normalized based on the gDNA controls. All primer and construct sequences are presented in Supplementary Table S1.

### Bioinformatic analyses

Raw ONT reads (fast5 format) were first basecalled using the Guppy software provided by Oxford Nanopore Technologies. We used Guppy version 6.4.2 on a GPU with the following configurations: --config rna_r9.4.1_70bps_hac.cfg --min_qscore 7 --records_per_fastq 10000000 --gpu_runners_per_device 8 --num_callers 1. The reads passing Guppy (fastq format) were then concatenated and aligned to the reference genome of *P. falciparum* 3D7 version 63 using minimap2 [38] and the following options: -ax map-ont -t 2. We then used samtools [39] to sort, index, and generate a bam output file for further data analysis. The data was visualized using IGV version 2.16.0 (Integrative Genome Viewer [40]). All further data analysis was performed using custom scripts.

The total number of reads at different categories (protein-coding genes, rRNAs, annotated ncRNAs, and pseudogenes) were counted based on reads that overlap ≥ 80% of the lengths of the read with one of these annotated features using custom scripts. Reads mapping outside of one of these annotated features or overlapping less than 80% were classified as “other”. To distinguish between the different 28S rRNA types, mapped reads overlapping ≥ 80% of PF3D7_0532000 or PF3D7_0726000 were classified as A-type, reads overlapping ≥ 80% of PF3D7_0112700 as S1-type, and reads overlapping ≥ 80% of PF3D7_1148640 or PF3D7_1371300 as S2-type.

To analyze the degradation at A-type 18S rRNA (PF3D7_0725600 and PF3D7_0531600) and 28S rRNA (PF3D7_0532000 and PF3D7_0726000), the numbers of reads of a given length overlapping ≥ 80% of one specific rRNA were counted, separately for each rRNA, and used for determining the 25%, 50%, and 75% quantiles. For both the read length and calculating the percentiles of rRNAs, 20 bp were added to each rRNA’s start and end position. For plotting the read coverage at 18S rRNA (PF3D7_0725600 and PF3D7_0531600) and 28S rRNA (PF3D7_0532000 and PF3D7_0726000) the number of reads overlapping each position was counted. To examine whether stretches of adenosines (As) in the genomic sequence was correlated with the starts of the ONT reads, As were counted across 20 bp windows sliding by 10 bp using the mpileup file generated with samtools for chromosome 7 and converted into a bedgraph file using custom scripts.

For analyzing the distribution of read lengths across all protein-coding genes, reads overlapping ≥ 80% of annotated protein-coding genes were counted and split by read length before being plotted with R Studio. In addition, the number of reads overlapping ≥ 80% of each of the 5,318 annotated protein-coding genes was calculated and this table was used for principal component analysis of the NF54 and Dd2 samples in R Studio. To gain information about the most abundant genes, we determined the minimum normalized read counts in each sample of the NF54 and Dd2 data sets, separately. For the NF54 samples, we included rings, trophozoites, schizonts, and stage V gametocytes from replicate #1. For Dd2 samples, the tightly synchronized gametocyte time course including all asexual and sexual stages was included. For the top 20 most abundant genes (excluding mitochondria and apicoplast genes), the number of reads of a specific read length was calculated for each gene separately.

## Results

### Direct RNA sequencing enables characterization of the NF54 *P. falciparum* transcriptome

We used Oxford Nanopore Direct RNA sequencing to analyze RNA from blood-stage *P. falciparum* NF54 parasites, including rings, trophozoites, schizonts and stage V gametocytes. We generated between 800,000 and 1,214,218 total reads from each stage, of which 689,756 - 1,067,087 reads passed base-calling (84 - 89%) and 593,185 - 969,642 reads mapped to the reference genome *Pf.* 3D7 (58 - 91%) (Supplemental Table S2). Between 50 and 70% of the reads overlapped with protein-coding genes enabling the characterization of transcripts from 3,589-4,198 annotated genes (Supplemental Table S2).

### Robust determination of the type of rRNA expressed at different developmental stages

ONT Direct RNA sequencing relies on the ligation of an oligo-dT/motor protein complex to the polyadenylated end of RNA molecules and therefore primarily targets mRNA molecules. However, due to their abundance and the relatively low specificity of the oligo-dT (see below), a high number of reads (between 13 and 22% of aligned reads) mapped to annotated ribosomal RNA (rRNAs), allowing detailed analyses of rRNAs (Supplemental Table S2). Since the ONT data is made of long reads, it is well-suited to differentiate the origin of the rRNAs that are highly similar to each other and, therefore, difficult to distinguish using short-read RNA-seq. In all asexual stages and stage V gametocytes of NF54 parasites, >99.7% of all rRNA reads unambiguously mapped to A-type rRNAs on chromosomes 5 and 7, with less than 0.3% mapping to S1 or S2 types (Figure 1A). We also used ONT Direct RNA sequencing to analyze RNA extracted from NF54 salivary gland sporozoites. Despite less than 2% of the reads mapping to the *Plasmodium* genome due to high contamination with mosquito RNA (Supplemental Table S2) [41, 42], 6,096 reads mapped to the different rRNA genes in roughly equal proportion: 38% of those reads aligned to A-type rRNAs, 37% to S1-type, and 24% S2- type rRNAs (Figure 1A, Supplemental Table S3). To validate these observations and confirm that ONT Direct RNA sequencing provided robust information on rRNAs despite technically targeting polyadenylated molecules, we designed type-specific primers for qRT-PCR and rigorously assessed the expression of the different rRNAs (see Materials and Method for details). In this experiment, we also included RNA extracted from mosquitoes fed on blood containing mature NF54 gametocytes and isolated 2-, 24-, and 72 hours post-infection, as well as RNA from the gametocyte culture which was used for membrane feeding (Figure 1B). Overall, these data confirmed the findings from ONT Direct RNA sequencing and showed that A-type rRNAs largely predominate in asexual blood stage parasites and gametocytes and persist after transmission to the mosquito, although S1- and S2 rRNAs gradually increase in proportion over time (Figure 1B). (Data for blood-stage parasites of the Dd2 strain are presented on Figure S1).

**Figure 1.**
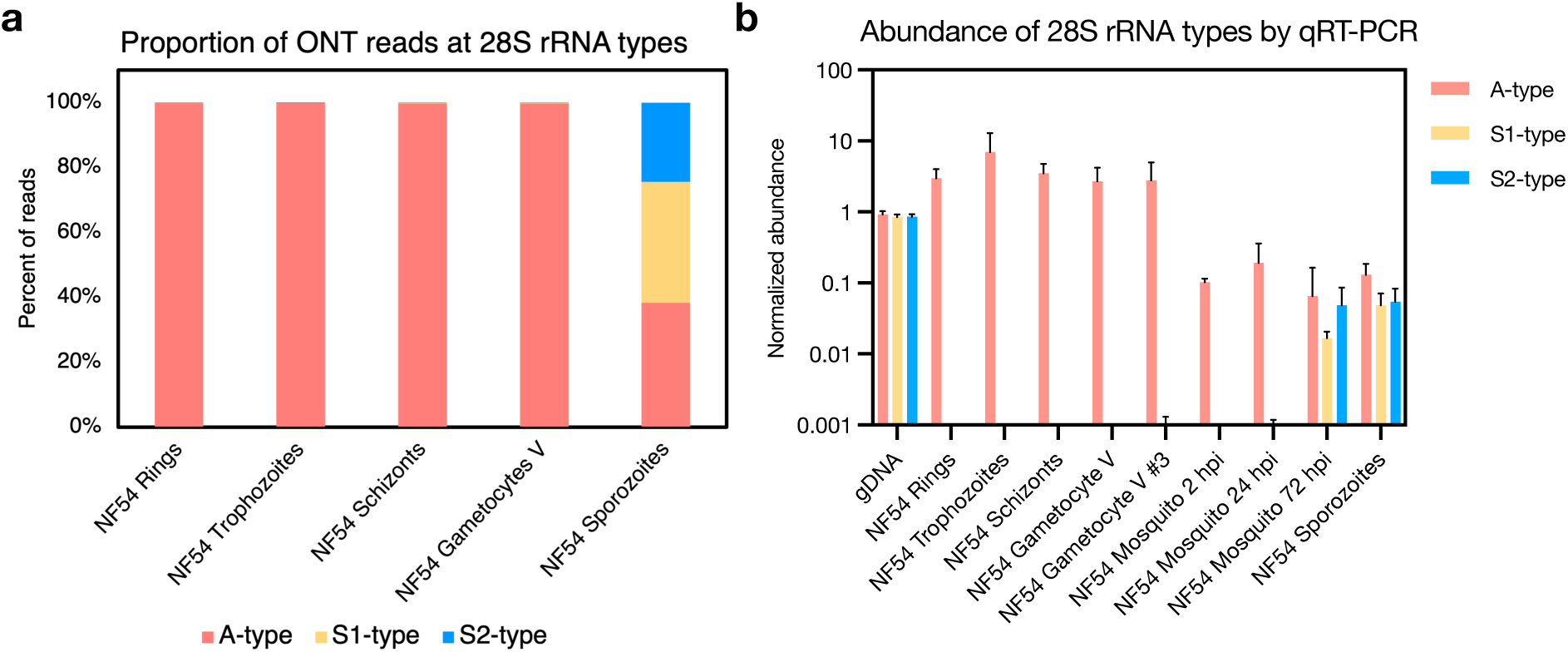
Proportion and abundance of A-type, S1-type and S2-type rRNAs in *P. falciparum* NF54 parasites. **(A)** Relative proportion of the three rRNA types in NF54 asexual stages (rings, trophozoites & schizonts), stage V gametocytes and salivary gland sporozoites based on ONT Direct RNA Sequencing. **(B)** Relative abundance of the three rRNA types based on qRT-PCR in NF54 rings, trophozoites, schizonts, stage V gametocytes and salivary gland sporozoites, as well as in an independent second stage V gametocyte culture used to feed mosquitoes (replicate #3) and in RNA extracted from mosquitoes 2-, 24- and 72 hours post infection (hpi). The raw qRT-PCR data was normalized using an exponential decay equation based on genomic DNA controls. The mean with the standard derivation is displayed (n = 3).

### rRNA degradation in *P. falciparum* NF54 gametocytes

We then examined variations in sequence coverage along the 18S and 28S rRNA genes. ONT sequencing starts from the oligo-dT adapter complexed with a motor enzyme and progresses as the RNA molecule passes through the nanopore. The sequencing, therefore, starts from the 3’ end, leading to the highest coverage there, and the coverage typically decreases along the sequence (due to sequencing of fragmented molecules and loss of sequencing signal), leading to a decrease in coverage from 3’ to 5’. While rRNA transcripts are not polyadenylated, the coverage observed in rings, trophozoites, and schizonts followed the same pattern for these genes with, especially for the 28S rRNA, a gradual decrease in coverage from 3’ to 5’ (Figure 2C). However, we also observed peaks of coverage in the middle of the rRNA genes (*e.g.*, around position 1,084,900 bp for the 18S rRNA gene and 1,088,000 bp for the 28S rRNA gene). These peaks coincide with stretches of adenosines (Figure 2A, 2C & S3) and likely represent sequencing of fragmented rRNAs ending with As that, although less abundant, are likely to be better captured and therefore disproportionally represented compared to full length rRNAs.

**Figure 2.**
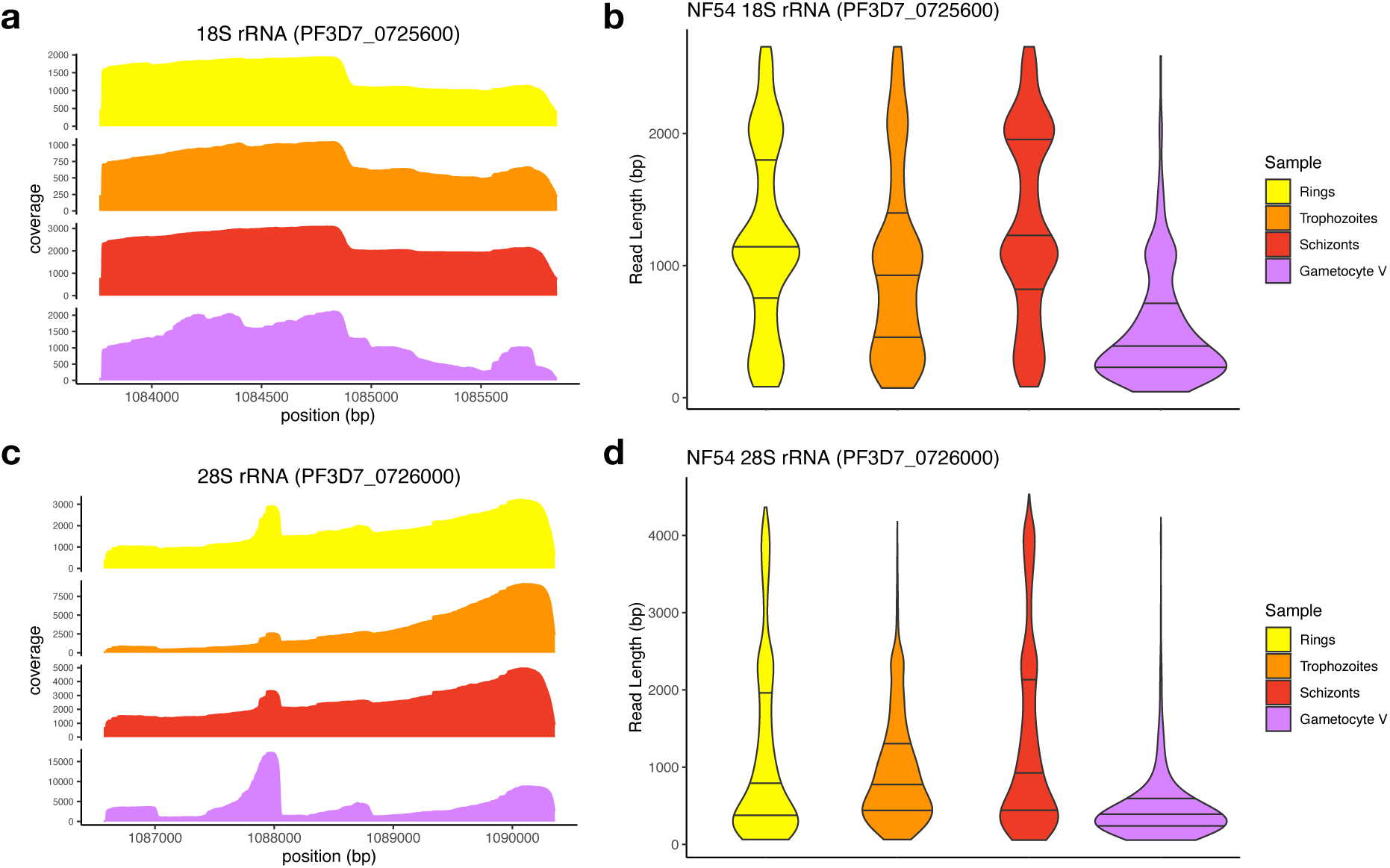
Analysis of ONT reads derived from the A-type 18S and 28S rRNAs in *P. falciparum* NF54 parasites. The figures show the coverage **(A, C)** and read length distributions **(B, D)** of the ONT Direct RNA Sequencing reads aligned to the 18S **(A, B)** and 28S rRNA genes **(A, D)** on chromosome 7 (PF3D7_0725600 and PF3D7_0726000, respectively). The data for ring-stage parasites is shown in yellow, trophozoites in orange, schizonts in red and stage V gametocytes in purple.

Surprisingly, the variability in coverage along the 18S and 28S rRNA genes appeared greater in stage V gametocytes than in asexual stages, possibly suggesting a higher proportion of degraded rRNAs in gametocytes (Figure 2A, 2C). To further investigate this pattern, we examined the lengths of the reads mapped to the 18S and 28S rRNA genes in asexual stages and stage V gametocytes. While many reads covered the entire length of the rRNA genes in asexual parasites, in stage V gametocytes most reads mapped to rRNA genes were very short: the median 18S rRNA read lengths for asexual stages were 911 – 1,122 bp for asexual parasites versus 344 bp for stage V gametocytes (p < 5.45×10^-15^) and 617 - 803 bp versus 342 bp for 28S rRNA (p < 3.89×10^-11^) (Figure 2B,D, Table 1). A similar pattern was observed for the A-type rRNA genes on chromosome 5 (Figure S2). Overall, these patterns suggested that rRNAs were more degraded in stage V gametocytes than asexual parasites.

**Table 1:** Summary of the ONT read lengths for A-type 18S rRNA (PF3D7_0725600) and 28S rRNA (PF3D7_0726000) from NF54 asexual stages and stage V gametocytes (including trophozoites and stage V gametocytes sequenced after *in vitro* polyadenylation). The table shows the median length (and the 25-75%iles in brackets).

| Sample | 18S rRNA read length (in bp) | 28S rRNA read length (in bp) |
| --- | --- | --- |
| NF54 Rings | 1,108 (657 – 1,885) | 617 (247 – 1,861) |
| NF54 Trophozoites | 911 (372 – 1,376) | 734 (418 – 1,263) |
| NF54 Schizont | 1,122 (793 – 2,012) | 803 (376 – 2,030) |
| NF54 Gametocyte V | 344 (206 – 663) | 343 (213 - 506) |
| NF54 Trophozoites +polyA | 1,568 (655 – 2,001) | 1,442 (664 – 3,263) |
| NF54 Gametocyte V +polyA | 218 (164 – 271) | 242 (181 - 339) |

### RNAs from protein-coding genes appear to be equally preserved in gametocytes and asexual parasites

To evaluate whether the putative degradation of rRNAs observed in stage V gametocytes was specific to rRNA genes or the result of general degradation of RNAs in gametocytes, we analyzed the length of reads mapped to protein-coding genes. While different genes are expressed at different stages, the overall distributions of transcript length inferred from the ONT reads were comparable: the median read length for protein-coding mRNAs in rings, trophozoites, and schizonts was 1,247 bp, 1,243 bp, and 1,313 bp, respectively, while the median read length for protein-coding genes in stage V gametocytes was 1,104 bp, only 10% shorter than in asexual parasites on average compared to a >50% differences in rRNA lengths (Table 1, Figure 3A, Table S3). This observation suggests that the RNA molecules are, overall, similarly preserved in the different samples. We also examined the read lengths obtained from each stage for genes that are highly expressed in all blood-stage parasites (Table S4) and observed no systematic differences in read length generated from asexual stages and gametocytes (Figure 3B-E, Figure S4). Overall, these analyses indicated that there were no major differences in mRNA degradation between asexual parasites and gametocytes and that the patterns of degradation observed in stage V gametocyte rRNAs appeared to be specific to rRNA genes.

**Figure 3.**
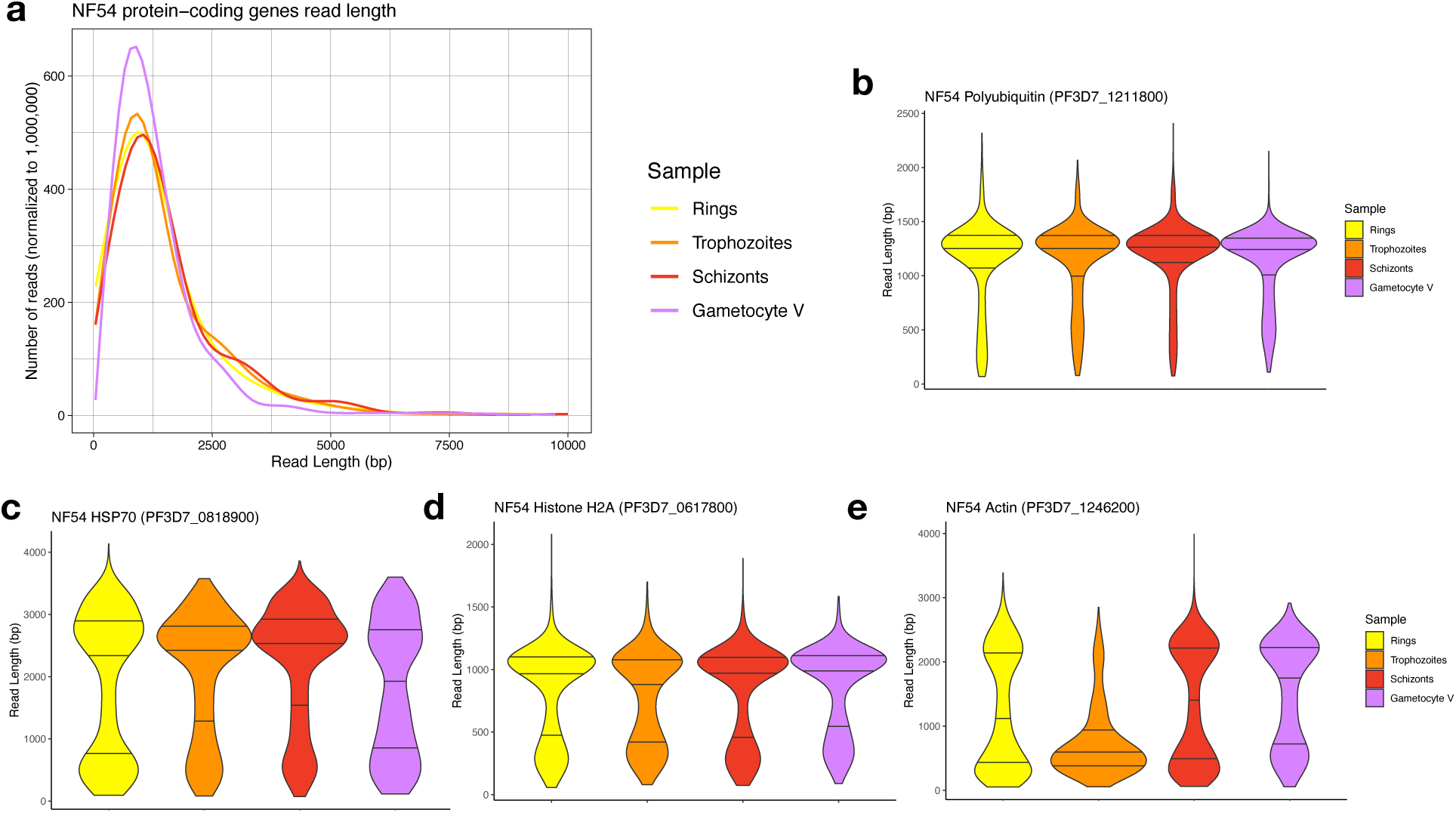
Analysis of the length of ONT reads derived from protein-coding genes. **(A)** Distribution of the lengths of reads (x-axis, in bp) mapped to protein-coding genes for NF54 ring-stage parasites (yellow), trophozoites (orange), schizonts (red) and stage V gametocytes (purple). **(B-E)** Distribution of the read length mapped to four highly expressed genes: *polyubiquitin* (PF3D7_1211800, **B**), *hsp70* (PF3D7_0818900, **C**), *histone H2A* (PF3D7_0617800, **D**) and *actin* (PF3D7_1246200, **E**).

### *In vitro* polyadenylation confirms that rRNA molecules are shorter in gametocytes

ONT Direct RNA sequencing captures rRNA transcripts due to their high abundance and the presence of stretches of adenosines at the end of degraded molecules (Figure S3). However, this non-specific capture prevented us from obtaining a comprehensive picture of the extent of the degradation in the rRNA, both in terms of proportion of degraded molecules compared to full length rRNA transcripts, and of the range and diversity of degraded molecules. We therefore repeated ONT sequencing after performing *in vitro* polyadenylation on all RNA molecules extracted from trophozoites and stage V gametocytes (see Materials and Method for details).

In both samples, after *in vitro* polyadenylation, the proportion of reads mapping to rRNA genes increased (Table S2), reflecting the abundance of these molecules, but with striking differences in read length between the two stages. In trophozoites, the sequence coverage became more even, with a slow and monotonous decrease towards the 5’ end of the 18S and 28S rRNAs (Figure 4A, 4C). In addition, many reads corresponded to full-length transcripts with, for example, a median read length of 1,442 bp at the 28S rRNA on chromosome 7 (Figure 4D, Table 1). Overall, *in vitro* polyadenylation increased the length of the reads mapped to rRNA genes expressed in trophozoites, indicating that i) most of these transcripts were full-length molecules and ii) that sequencing without *in vitro* polyadenylation led to an over-representation of degraded transcripts. By contrast, after polyadenylation of stage V gametocyte RNAs, we observed an overwhelming abundance of very short rRNA reads (with a median read length of 242 bp for the 28S) and almost no full-length transcripts (Figure 4B, 4D, Table 1). The short reads in stage V gametocytes were also distributed throughout the entire 18S and 28S rRNA gene sequences with no clear pattern or peak, except for an accumulation of reads at the 5’ and 3’ end of the 28S rRNA gene (Figure 4A, 4C). This pattern is consistent with a role for exonucleases in the rRNA degradation rather than a sole cleavage by an endonuclease or, alternatively, for a cleavage at numerous sites distributed throughout the rRNA sequences. Thus, while the *in vitro* polyadenylation in trophozoites enabled sequencing full-length (non-polyadenylated) rRNA transcripts, the same experiments in stage V gametocytes confirmed that most rRNAs were highly degraded and that, proportionally, very few full-length rRNA transcripts were present in these parasites.

**Figure 4.**
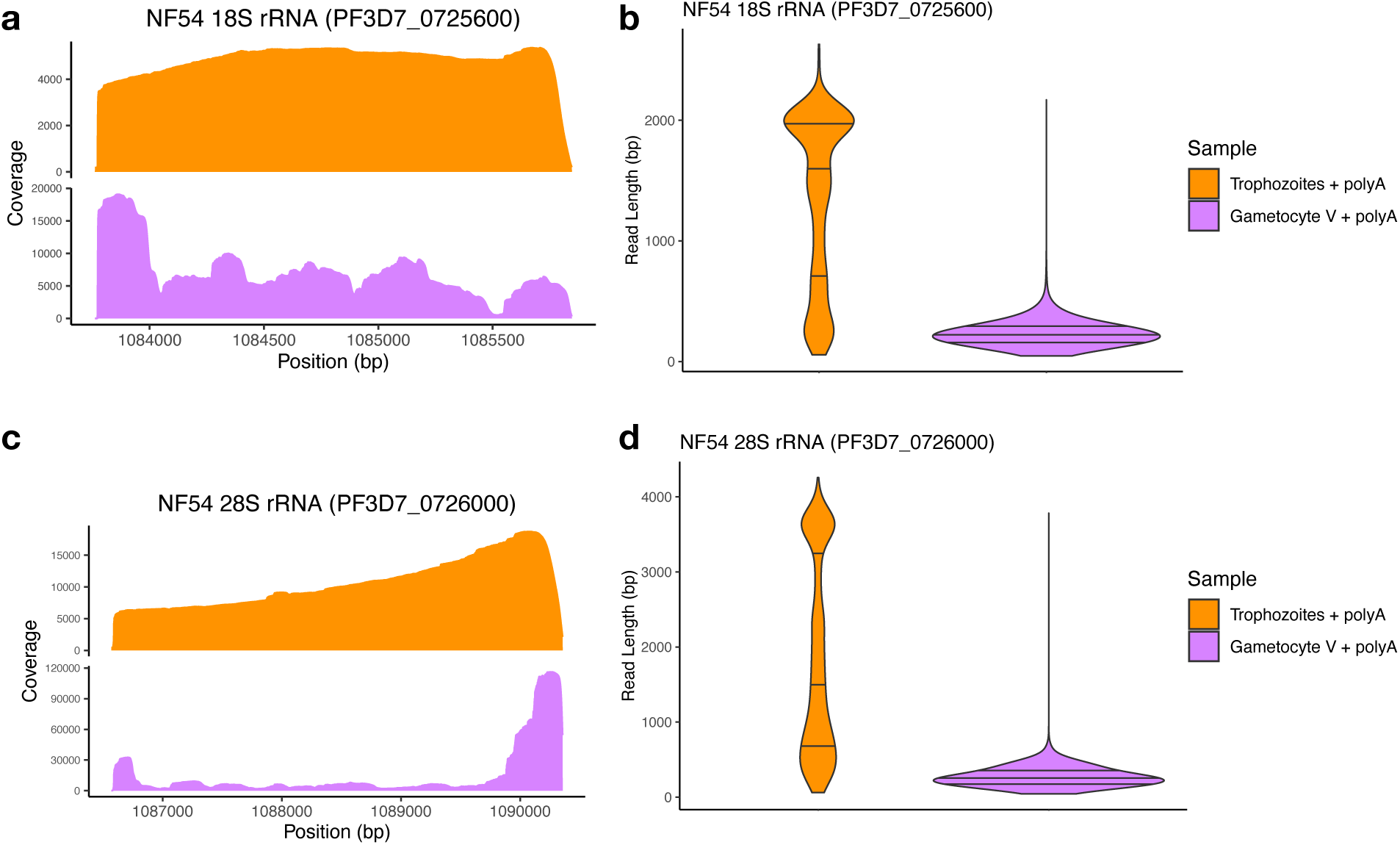
Coverage profile **(A, C)** and read length distribution **(B, D)** of Direct RNA sequencing ONT reads generated after *in vitro* polyadenylation of NF54 trophozoites (orange) and stage V gametocytes (purple) mapped to the A-type 18S and 28S rRNA on chromosome 7.

### Replication and determination of the timing of rRNA degradation during gametocytogenesis

To evaluate whether the rRNA degradation observed could have been artificially induced by the culture protocol, we sequenced RNA from an independent gametocyte culture of the same NF54 strain (see Materials and Method). Importantly, this second sample derived from cultures routinely used to feed mosquitoes and study parasite transmission at the Johns Hopkins Bloomberg School of Public Health. This second gametocyte sample also showed a highly degraded coverage profile of rRNAs, possibly of a greater extent than in the first sample (Figure S5).

To gain insight on the timing of rRNA degradation and ensure that the patterns observed were not specific to the NF54 strain of *P. falciparum*, we sequenced RNAs collected throughout gametocyte development of *P. falciparum* Dd2 parasites, including sexually committed rings and gametocytes stage I - V as well as asexual parasites (10-, 20-, 30- and 40 hpi). We generated 525,150 - 1,686,943 reads from each sample (Table S2), with 42 - 80% of reads overlapping protein-coding genes and 4 - 34% of the reads mapping to rRNA genes (Table S2). To ensure that similar developmental stages were compared in NF54 and Dd2 (and rule out technical outliers/artefacts), we performed a principal component analysis (PCA) based on the gene expression determined from each sample at protein-coding genes. As expected, asexual life cycle stages from NF54 and Dd2 clustered together, while the NF54 stage V gametocyte sample clustered with the Dd2 stage V (day 12 post sexual commitment) and V d16 (day 16 post sexual commitment) gametocyte samples (Figure 5A), highlighting the overall similarities between life cycle stages in both strains. We also analyzed the read length of reads mapped to protein-coding genes and found no major difference between the length of mRNAs sequenced from asexual stages and gametocytes (Figure 5B, Figure S7, Table S3), confirming the similar quality of the RNA samples from different stages.

**Figure 5.**
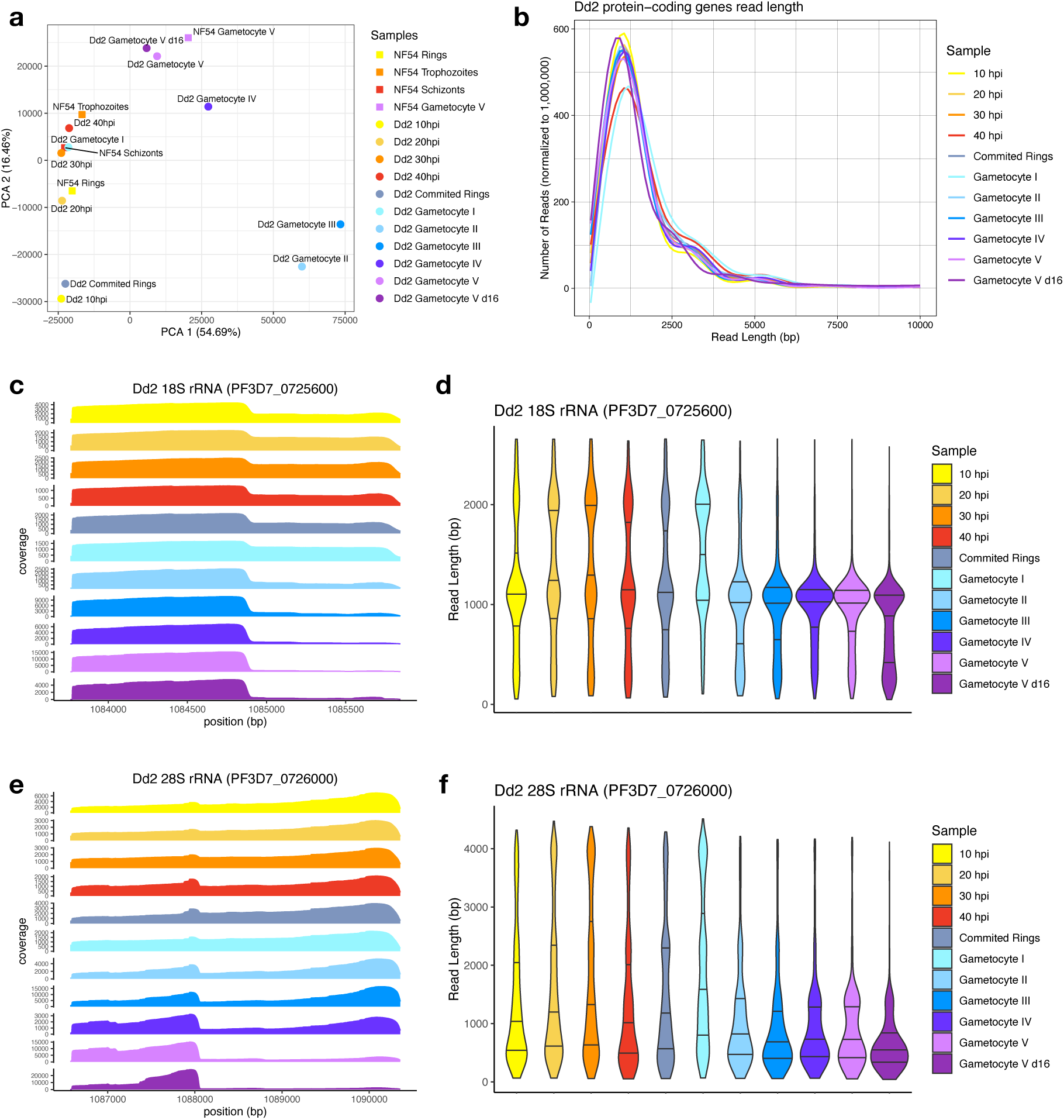
Analysis of ONT Direct RNA Sequencing generated from the Dd2 *P. falciparum* strain. **(A)** Principal component analysis of the gene expression profiles determined by ONT sequencing at protein-coding genes for NF54 (squares) and Dd2 (circles) asexual and sexual stages. Each point is colored according to the parasite stage: rings in yellow, trophozoites in orange, schizonts in red, sexually-committed rings in grey, and stage I to V gametocytes in blue to purple. **(B)** Distribution of the lengths of reads mapped to all protein-coding genes. **(C-F)** Coverage and read length distribution of the ONT Direct RNA-sequencing reads from Dd2 parasites mapped to the 18S **(C-D)** and 28S **(E-F)** rRNA genes.

The patterns of read coverage and read length at rRNA genes in Dd2 parasites were, overall, very similar to those observed for NF54. Again, the ONT reads mapped to rRNAs were significantly shorter in stage V gametocytes than in asexual parasites, consistent with a degradation of rRNAs in gametocytes (Figures 5C-F). As previously, we also sequenced RNAs from one early trophozoite sample (20 hpi) as well as stage V gametocytes from day 12 and day 16 after *in vitro* polyadenylation and the results were qualitatively similar to those observed in NF54 parasites, although the rRNAs degradation appeared less pronounced in Dd2 stage V gametocytes (Figure S6).

The collection of RNA from Dd2 parasites at different stages of their gametocytogenesis enabled us to narrow down the timing of the rRNA degradation. While the data for sexually committed rings and stage I gametocytes were consistent with the presence of full-length rRNAs at these stages, we observed an abrupt shift in stage II gametocytes that appeared to primarily contain degraded rRNA transcripts and this rRNA degradation continues over time, with even more fragmented rRNA in mature gametocytes on day 16 (Figure 5C-F).

## Discussion

The life cycle of *Plasmodium* parasites is characterized by complex gene regulation ensuring a rapid response to environmental changes, notably those occurring during the transmission of the parasites between the mammalian host and the insect vector. As *P. falciparum* progresses through its life cycle, it relies on three different types of rRNA, the A-, S1- and S2-type, that have been previously described to vary in expression between different life cycle stages [25]. The expression of these different rRNA types may allow for optimization of translation in different conditions (e.g., temperature or pH) or of different genes, or could be associated with different ribosome composition (e.g., ribosomal proteins) [28]. However, the molecular mechanisms underlying the regulation and turnover of these different rRNAs in *Plasmodium* is poorly understood, with most studies focusing exclusively on the abundance of rRNAs at different stages while the transcription of these genes is comparatively understudied and their degradation rarely considered [21–23, 26, 29, 30].

We have used Oxford Nanopore Direct RNA sequencing to analyze the dynamics of *P. falciparum* rRNA expression during the intraerythrocytic asexual development and over the time course of gametocytogenesis, as well as in mosquito salivary gland sporozoites. The long reads generated by ONT sequencing are particularly suited for this purpose as they enable us to robustly differentiate closely related nucleotide sequences, such as those of the different rRNA types. Our analysis confirmed that A-type rRNAs account for the vast majority of rRNAs expressed by asexual blood-stage parasites, as previously described in the literature [21, 29, 30]. More surprisingly, A-type rRNAs remain the dominant rRNAs throughout gametocytogenesis, up to and including stage V gametocytes. This observation may appear to contrast with previous literature describing the S1-type as the most actively transcribed rRNA in these stages [30]. In fact, in the tightly synchronized Dd2 gametocyte time course, we also observed the appearance of the S1-type rRNA, but with a very low read number in stage III gametocytes (Table S3). However, transcription of the S1-type rRNA in stage III gametocytes does not dramatically affect the relative abundance of rRNAs and the A-type rRNAs still account for the majority of all rRNAs (and this finding could also be reproduced by qRT-PCR, see e.g., Figure 1B). S-type rRNAs are thought to dominate during the development of *P. falciparum* in mosquitoes [22, 26, 30], but both ONT sequencing and qRT-PCR analyses showed that all types (A-, S1-, and S2-types) were present in roughly equal proportion in salivary gland sporozoites. Consistent with this observation, we noted an increase in the abundance of both S-type rRNAs in *P. falciparum* parasites collected from mosquitoes at 2-, 24- and 72 hours after an infected blood meal, but in a context where the A-type rRNAs persisted.

We then leveraged the ONT sequences to study the extent and dynamics of rRNA degradation throughout *P. falciparum* development. Many of the ONT Direct RNA sequencing reads capture the entire length of the transcripts, enabling examining the size distribution of RNA molecules from a given locus. This feature contrasts with traditional methods of RNA analyses, such as short read RNA-seq, RT-qPCRs or fluorescence *in situ* hybridization (FISH), that only examine a small region of the RNA sequence and do not easily allow differentiating full-length transcripts from fragmented molecules. Northern blots remain the gold standard for examining the length of various RNA molecules but i) are quite cumbersome experimentally and ii) require multiple carefully designed probes to obtain a specific and comprehensive perspective. The length of ONT reads mapped to rRNA genes, and their distribution throughout the genes indicated that, while the 18S and 28S rRNAs expressed in asexual parasites matched their annotations and supported intact, full-length transcripts, these rRNAs appeared to be fragmented in gametocytes, regardless of the *P. falciparum* strain examined or of the culture methods used for generating gametocytes. This degradation of rRNAs in gametocytes was not mirrored in protein-coding mRNAs that appeared equally preserved in gametocytes and asexual blood stage parasites, indicating that this degradation, and its underlying mechanism, was specific to rRNAs. Analysis of rRNAs throughout gametocytogenesis revealed that degradation of A-type rRNAs starts in stage II gametocytes and progresses over time, leading to a significant reduction in the numbers of full-length rRNAs (although the extent of this reduction may vary among stage V gametocyte samples, possibly reflecting small differences in their “maturity”). Recent studies in humans and other eukaryotes have shown that rRNAs can be processed into ribosomal RNA fragments (rRFs) by specific cleavage mediated through endonucleases [43]. These short rRFs (typically 15-40 nucleotide long) can then regulate transcription or translation of specific genes [43]. In stage V gametocytes, we did not see enrichment of specific fragments resulting from rRNA degradation (except at extremities of the rRNA molecules) but rather a general and even degradation of rRNAs without clear pattern. While we cannot exclude that further processing eliminates most degraded fragments to exclusively keep specific rRFs, the most parsimonious explanation of our data is that the rRNAs are degraded to reduce their abundance in gametocytes, not to generate specific regulatory molecules.

Despite their critical importance, there is relatively little known about the mechanisms underlying rRNA turnover. Degradation of rRNAs and turnover of ribosomes has mainly been studied in prokaryotes and in some model eukaryotic organisms, such as *Saccharomyces cerevisiae,* where this process is notably associated with ribosome assembly errors and environmental stress response [44–46] and involves various ribonucleases [46]. rRNA turnover and degradation of *Plasmodium* has not been studied in depth. One study in 1970 showed that chloroquine treatment could lead to the degradation of rRNA in *P. knowlesi* [47] and the 18S A-type rRNA has been shown to be cleaved within a functional region at the 3’ end of the molecule, possibly affecting ribosome functionality [22]. This latter study, showing that some of the A-type 18S rRNA molecules were cleaved in gametocytes (and further processed in zygote and ookinete) [22], is consistent with our observations although our data suggest that there may be numerous cleavage sites in the 18S and 28S rRNAs and/or that these molecules are further processed by exonucleases.

In bacteria, during slower growth phases when the demand for protein synthesis is reduced, cells produce fewer ribosomes (sometimes up to tenfold fewer than in fast-growing cells) and actively degrade unassembled ribosomal subunits, including rRNAs [44]. In this regard, the observation of *Plasmodium* rRNA degradation during gametocytogenesis is particularly intriguing, because mature gametocytes remain in a poised state waiting for ingestion by a mosquito to trigger their further development [3, 17] and their metabolic activity is reduced as is gene transcription and translation [7, 48]. However, upon transmission to the mosquito, the gametocytes need to reactivate rapidly to complete gametogenesis within a few minutes [8–10, 49]. One of the mechanisms underlying this quiescence and rapid reactivation is translational repression: mRNAs from genes required for the development in mosquito are stabilized by the DDX6-class RNA helicase DOZI [14] and stored in messenger ribonucleoprotein particles (mRNPs) [50, 51]. Upon transmission, the mRNAs are released from these granules and made accessible for translation. In addition, the Eukaryotic Initiation Factor 2 subunit alpha (eIF2α) is phosphorylated by eIK2 in gametocytes, leading to a global repression of translation, effectively reducing overall protein synthesis [52]. In this context, it is tantalizing to speculate that the specific degradation of rRNAs during the late stages of gametocytogenesis might not only be important for the (partial) rRNA switch occurring during transmission but also play a key role in regulating translational repression and quiescence. Thus, rRNA degradation might occur as a consequence of the overall decrease in translation as more ribosomes become idle and are disassembled, leading to increased accessibility of putative cleavage sites as described in bacteria [44, 46]. Alternatively, one could speculate that the active degradation of rRNAs during gametocytogenesis (e.g., by expression of specific nucleases) could contribute to translational repression by reducing the number of ribosomes and lowering the translational capacity of these parasites.

## Acknowledgements

This study was supported by awards from the National Institutes of Health (R01AI172827 to DS, U19AI110820 to DS and JCDH, R01AI132359 to PS, and R03AI180804 to SK), from the Deutsche Forschungsgemeinschaft (DFG, German Research Foundation - 549182383 to JG), from Bloomberg Philanthropies (to PS, SK, AT), and T32-GM125592 to RS. We would like to thank the insectary and parasitology cores at the Johns Hopkins Malaria Research Institute. The funders had no role in study design, data collection and analysis, decision to publish, or preparation of the manuscript.

## Data and code availability

All sequence data generated in this study have been deposited in the National Center for Biotechnology Information (NCBI) Sequence Read Archive under the BioProject ID XXXX. Custom scripts are available at https://github.com/jgruenebast/rRNA_degradation.

## Notes

### Competing Interest Statement

The authors have declared no competing interest.

