## Supplemental Figures for "Degradation of ribosomal RNA during *Plasmodium falciparum* gametocytogenesis"

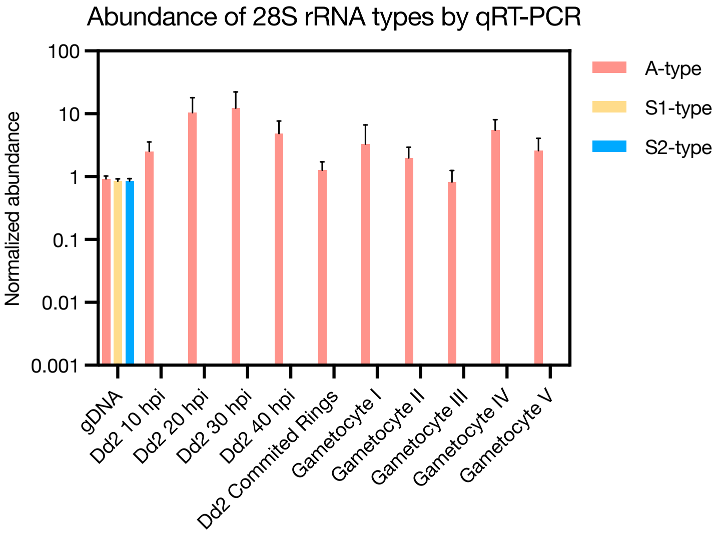


**Figure S1:** Relative abundance of the three rRNA types in Dd2 asexual and sexual parasites based on qRT-PCR. The mean with standard derivations is shown, n = 3.

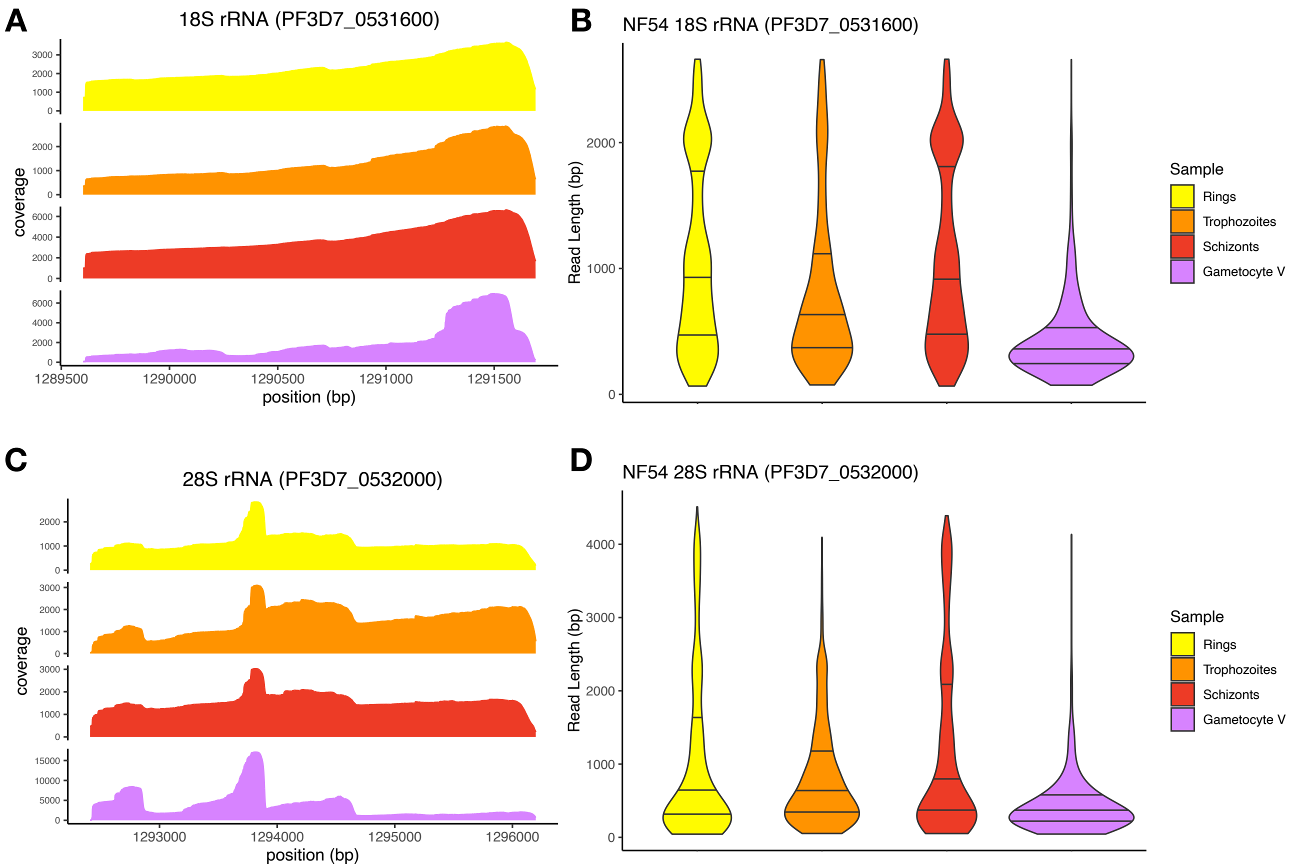


**Figure S2:** Coverage profile and read length distribution of ONT Direct RNA Sequencing reads from NF54 rings, trophozoites, schizonts and stage V gametocytes mapped to the 18S (A, B) and 28S rRNA gene (C, D) on chromosome 5.

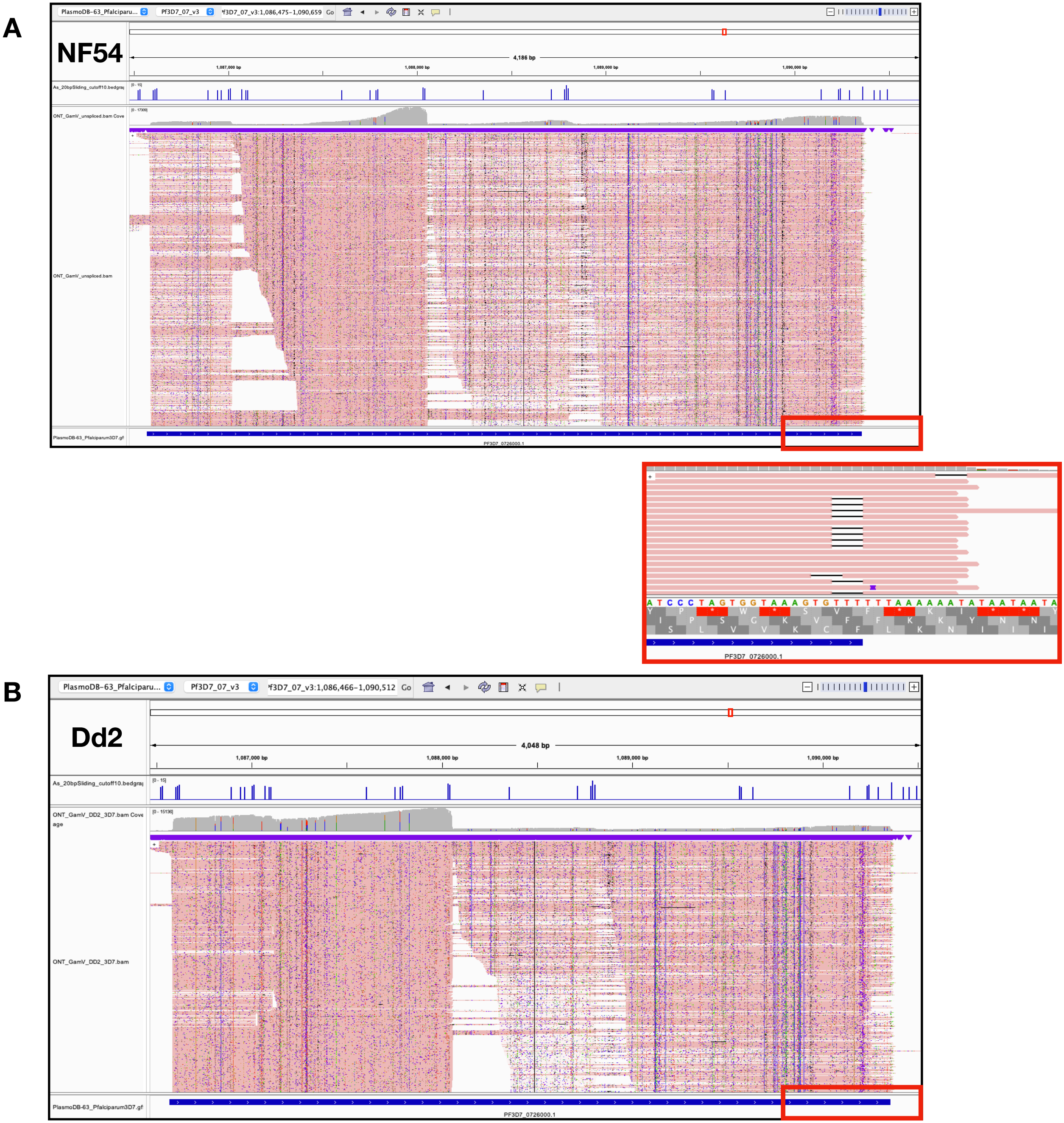


**Figure S3:** IGV screenshots showing the coverage (grey) of ONT NF54 stage V gametocyte reads (in pink) **(A)** and Dd2 stage V gametocytes reads **(B)** mapped to the 28S rRNA gene on chromosome 7 (PF3D7_0726000). The blue tick marks above the coverage tracks indicate the numbers (y-axis) of consecutive adenines in 20 bp sliding windows of the genome sequence (only windows with more than 10 adenines are displayed). The red box zooms in on the end of the 28S rRNA and highlights the stretches of adenosines.

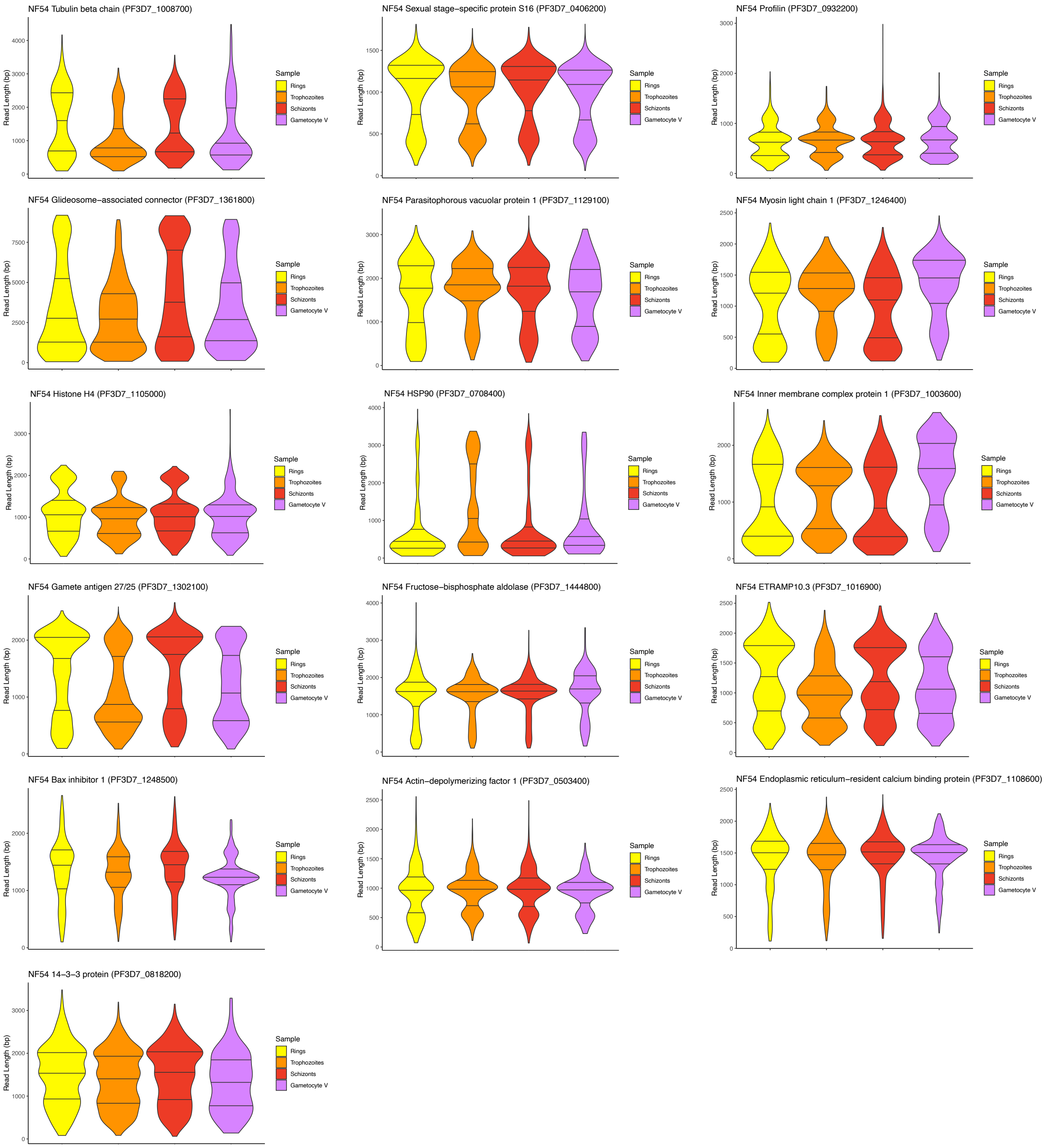


**Figure S4:** Read length distribution at the remaining 16 most abundantly expressed protein-coding genes in NF54 parasites.

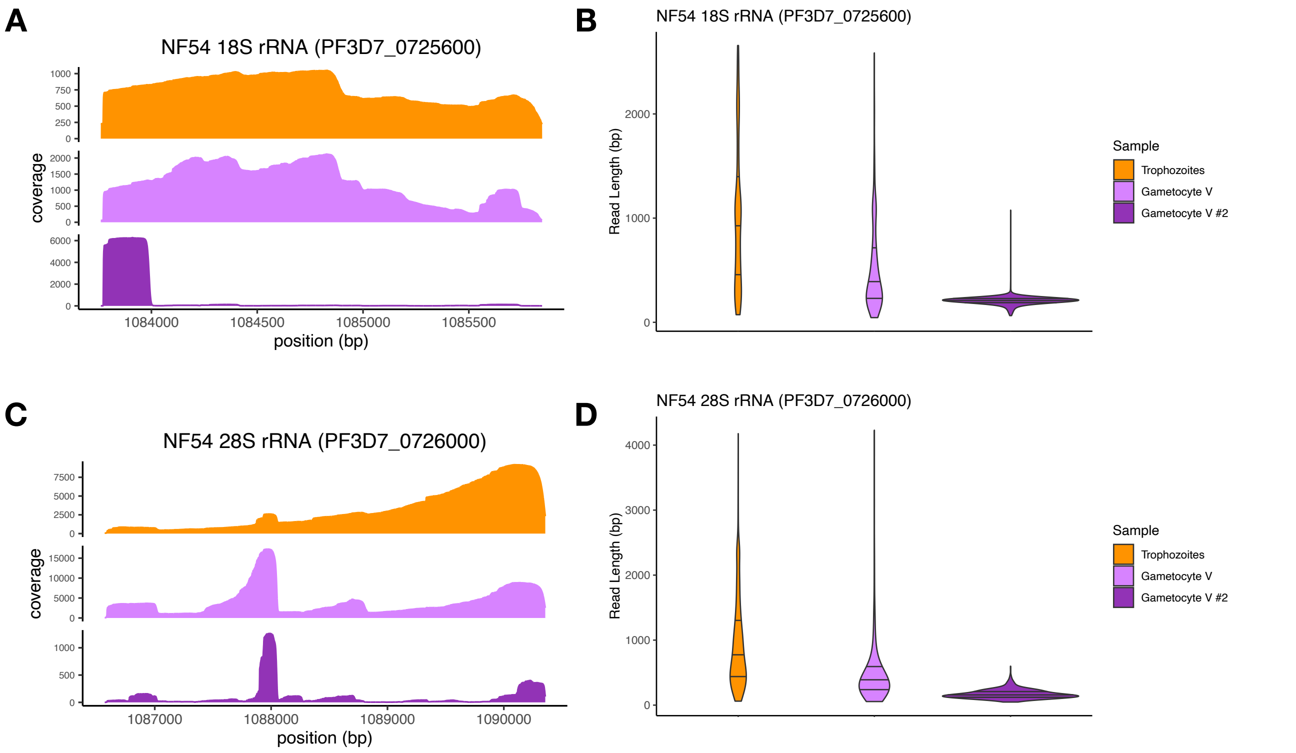


**Figure S5:** Coverage profile and read length distribution from ONT direct RNA sequencing reads stage V gametocytes replicate #2 (dark purple) compared to NF54 trophozoites and stage V gametocyte replicate #1. (A-B) 18S rRNA on chr. 7, (C-D) 28S rRNA on chr. 7.

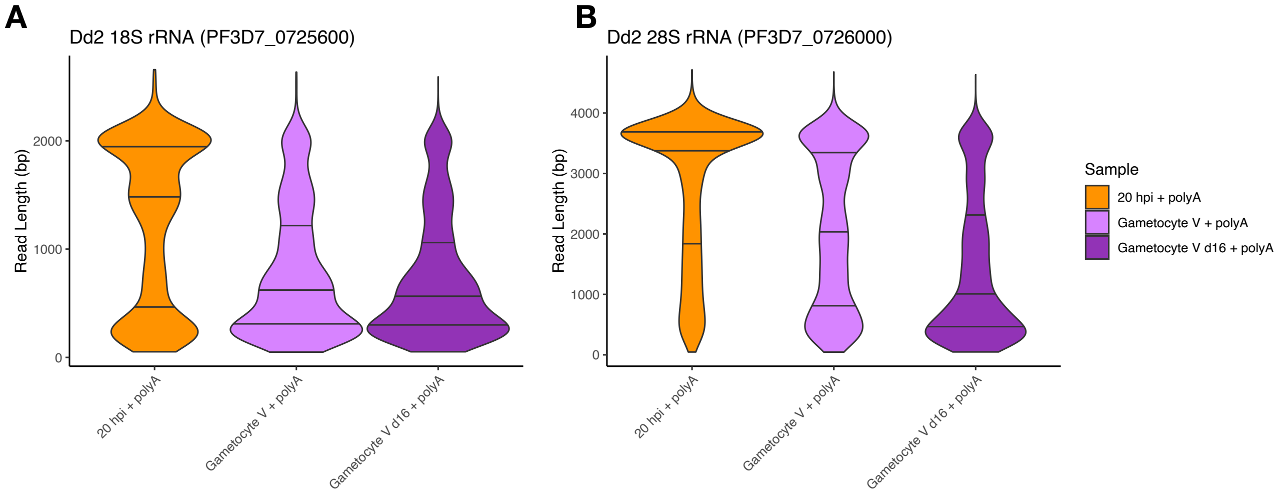


**Figure S6:** Read length distribution of ONT direct RNA sequencing after *in vitro* polyadenylation of *P. falciparum* Dd2 trophozoites, stage V gametocytes and gametocytes stage V day 16. (A) 18S rRNA on chr. 7, (B) 28S rRNA on chr. 7.

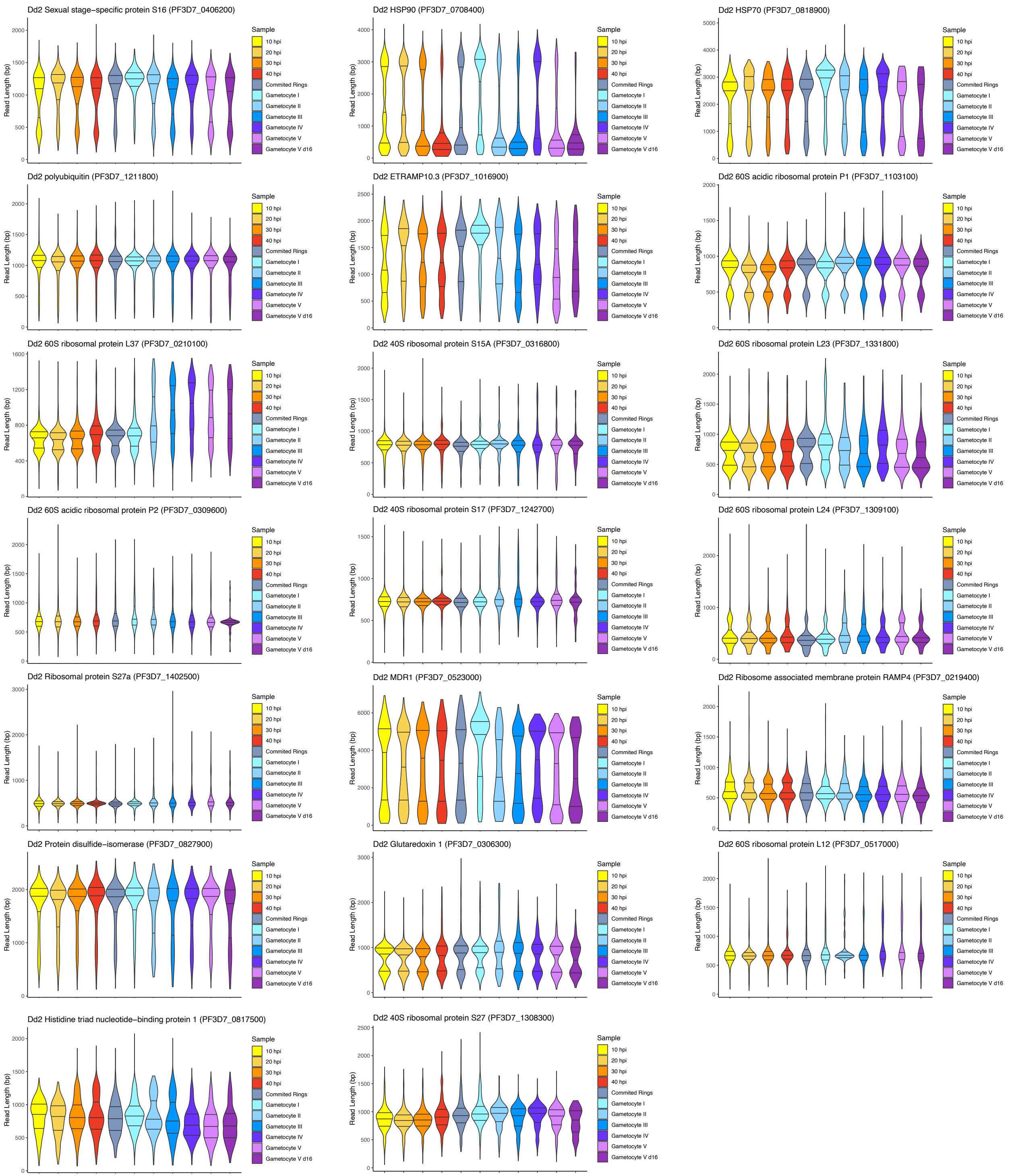


**Figure S7:** Read length distribution at the top 20 most abundant protein-coding genes in *Pf.* Dd2 parasites.
